# Neural signatures of predictive coding underlying the acquisition of incidental sensory associations

**DOI:** 10.1101/2025.05.16.654429

**Authors:** Antonino Greco, Clara Rastelli, Leonardo Bonetti, Christoph Braun, Andrea Caria

## Abstract

A longstanding question in cognitive science is whether the human brain learns sensory regularities that are irrelevant to ongoing behavior, a phenomenon known as incidental associative learning. Here, we provide evidence at the single subject level that humans indeed acquired such incidental associations and reveal their neural signatures by combining Electroencephalography (EEG), multivariate decoding and computational modeling. We found robust encoding of incidental predictions and prediction errors underlying associative learning, consistent with predictive coding theories. Prediction errors were modulated by epistemic uncertainty, incidental associations persisted in memory over days and generalized beyond stimulus-stimulus pairings to encompass broader sensory events, including the absence of stimulation. Together, our results challenge traditional assumptions that associative learning requires behavioral relevance or physical co-occurrence and point to uncertainty minimization about environmental state transitions as a general objective of brain function. This intrinsic drive may reflect a fundamental computational principle supporting the formation of internal predictive models that guide perception, learning, and adaptive behavior.

## Introduction

Biological systems continuously face the challenge of adapting to dynamic and ever-changing environments. To navigate these uncertainties, it is crucial to anticipate future events, recognizing cues that may indicate potential risks or rewards. The human brain is especially adept at extracting patterns from sensory information, allowing individuals to forecast changes in their surroundings and adjust their behavior accordingly. This predictive ability forms the foundation of many higher-order cognitive functions, supporting processes as diverse as perception, language comprehension, and flexible decision-making (*1*– *3*).

At the heart of the brain’s ability to adapt to its environment lies associative learning, the process by which relationships between sensory events are encoded and stored in memory (*4*–*9*). Since the earliest studies of classical (*10*–*13*) and instrumental conditioning (*14*–*17*), associative learning has been recognized as a fundamental building block of cognition (*18*–*20*). Building on this principle, contemporary research has shown that the brain also tracks complex patterns and regularities across streams of sensory input, even when these patterns are subtle or probabilistic. This ability, often referred to as statistical learning (*21*– *25*), highlights how associative processes extend beyond simple pairings to encompass the extraction of higher-order structures from the environment.

Predictive coding theory proposes that associative learning is not merely a passive recording of co-occurrences, but rather an active, predictive process (*26*–*36*). According to predictive coding theoretical framework, the brain constructs internal models that anticipate incoming sensory information by predicting future observations (*37*– *41*), and continuously updates these models based on the mismatch between expected and observed inputs, known as prediction errors (*42*–*53*). In this view, associative learning arises through the minimization of prediction errors, allowing organisms to efficiently model and navigate their environment (*54*–*56*).

While much of what we know about learning derives from studies in which humans are explicitly instructed to acquire new information, real-world environments rarely offer such structured and supervised opportunities. This realization has led to a growing interest in the distinction between explicit and implicit (associative or statistical) learning (*57*– *63*). Explicit learning is characterized by deliberate efforts to acquire knowledge, typically guided by task demands and accompanied by metacognitive processes. In contrast, implicit learning occurs without awareness of what has been learned, yet it typically concerns information that remains behaviorally relevant to the task at hand (*64*–*67*).

Within the domain of implicit learning, a particularly intriguing and debated phenomenon is incidental learning (*68*–*71*), the acquisition of sensory associations that are entirely task-irrelevant. In incidental learning paradigms, participants engage in an overt task unrelated to the regularities being embedded in the environment, such that the learned associations neither facilitate nor interfere with task performance. This crucial difference, the absence of behavioral relevance, distinguishes incidental learning from other forms of implicit learning and raises fundamental questions about the automaticity and scope of the brain’s learning mechanisms.

However, the existence and nature of incidental learning remain subjects of intense debate. A central question is whether the brain, in its drive for efficiency (*72*), actively minimizes energy expenditure by discarding predictions about task-irrelevant sensory events, or whether it instead preserves predictable regularities even when they carry no immediate behavioral relevance, maintaining them as a latent resource for potential future use. From an energetic standpoint, continuously tracking irrelevant associations would seem costly and unnecessary. Yet from an adaptive perspective, retaining hidden regularities could confer significant advantages in a dynamic environment where priorities can shift unexpectedly.

Empirical findings on this issue have been mixed. Some studies report that the brain selectively learns only behaviorally relevant sensory associations (*73*–*78*), showing little to no evidence of association formation when regularities are irrelevant to current goals. Others, however, demonstrate that sensory associations can be acquired incidentally (*79*–*86*), typically without awareness. Several factors may contribute to these discrepancies. On one hand, incidental learning may be inherently slower or less efficient than learning of task-relevant information, potentially requiring longer exposure durations than those typically employed in prior experimental paradigms. On the other hand, studies reporting incidental learning may be influenced by subtle forms of covert task relevance, whereby the associations being acquired are not entirely irrelevant to participants’ behavior or attentional priorities. Disentangling true incidental learning from these confounding factors remains a major challenge for understanding the breadth and flexibility of the brain’s predictive mechanisms.

Here, we sought to directly address this issue by combining electroencephalographic (EEG) recordings with multivariate decoding and computational modeling using an incidental learning paradigm. Human participants performed a target detection task while being exposed to a stream of auditory and visual distractor stimuli whose statistical structure was manipulated unbeknownst to them and was entirely irrelevant to task performance. Crucially, the experiment spanned six sessions over two weeks, providing extended exposure to the sensory regularities and allowing us to probe incidental learning under conditions in which, if present, it could reliably be observed.

Our investigation was guided by several key questions. First, we asked whether participants incidentally acquired sensory associations despite their lack of task relevance, and what neural mechanisms underpinned this learning. Second, if incidental learning happened, what were the neural signatures of predictions and prediction errors during the acquisition of incidental sensory associations? Third, motivated by the Bayesian brain hypothesis (*28, 30, 87*), we tested whether participants incorporated epistemic uncertainty to modulate the weighting of prediction errors over time. Fourth, we investigated whether the learned associations were retained across days, indicating memory consolidation, or whether they were relearned anew at each experimental session. Finally, our experimental design allowed us to test whether incidental predictive coding extends beyond direct stimulus-stimulus associations to encompass general sensory events (*6, 68, 88*–*91*), including periods of “sensory silence”. By addressing these questions, we aimed to provide a comprehensive characterization of how predictive coding mechanisms operate during the acquisition of incidental sensory associations and uncover their neural signatures.

## Results

We instructed human participants to perform a target detection task by being exposed to a stream of audio and visual stimuli. The task consisted of pressing a button upon detecting a target stimulus, while ignoring the distractor stimuli (Fig. 1A). Unbeknownst to the participants, we manipulated the transition probability between the distractor events. This allowed us to investigate implicit associative learning, defined as the acquisition of task-irrelevant or incidental sensory associations. Thus, we focused our investigation solely on the distractors and the associations among them. Importantly, none of the participants reported awareness of the manipulated associations between the distractor events, validating our experimental design. Distractor sensory events included pure auditory tones (A_1_ and A_2_) and visual Gabor patches (V_1_ and V_2_), including their absence (A_0_ and V_0_).

**Fig. 1.**
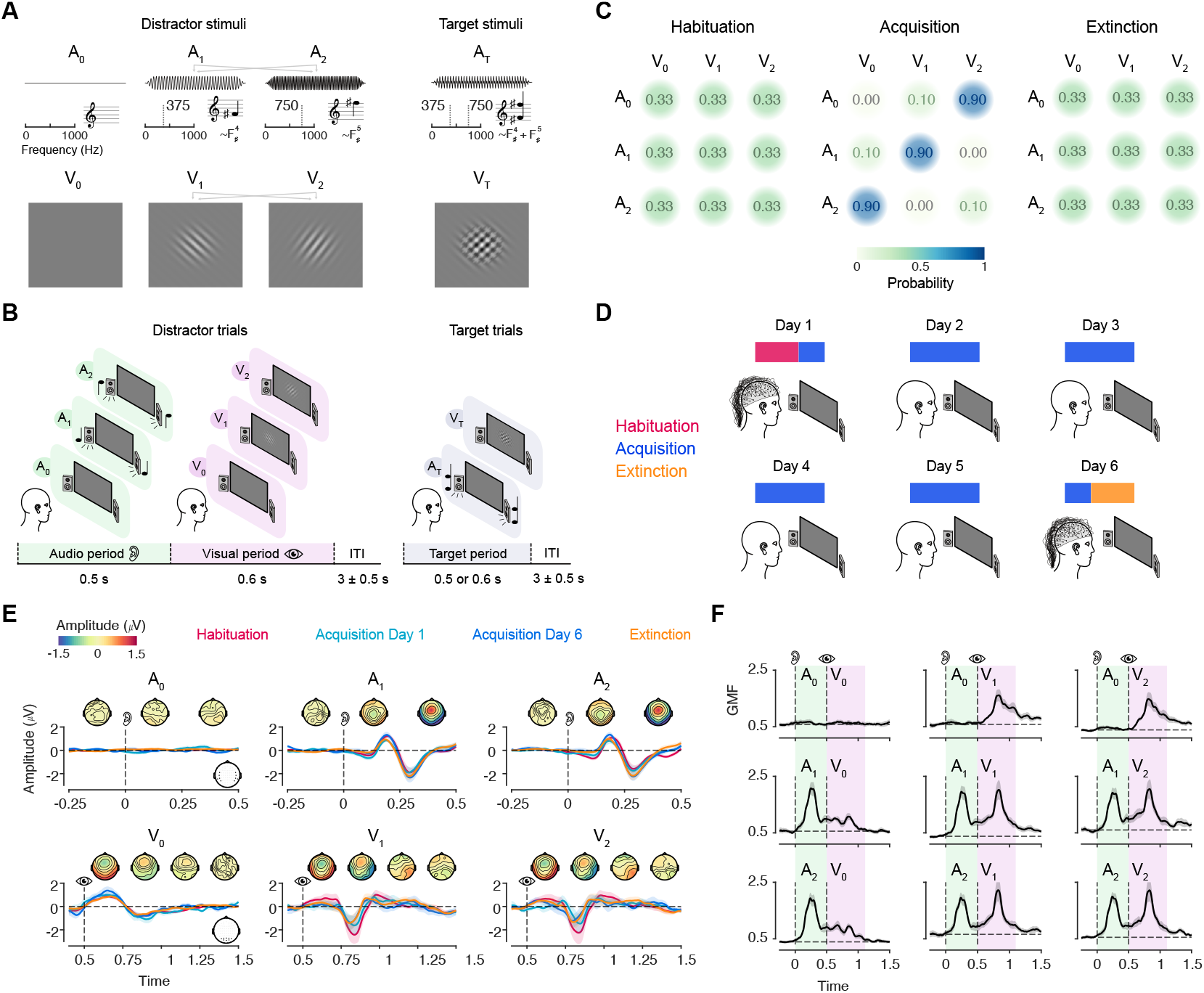
Experimental design and evoked responses to audio and visual stimuli. Participants were instructed to perform a target detection task by being exposed to a stream of audio and visual stimuli. The task consisted of pressing a button upon detecting a target stimulus, while ignoring the distractor stimuli. **A**. The distractor stimuli were two auditory stimuli (A_1_ and A_2_), low and high frequency pure tones of 375 Hz and 750 Hz (roughly corresponding to a fourth and fifth octave F_#_ musical note) and two visual stimuli (V_1_ and V_2_), Gabor patches with 45° and 135° orientation. The arrows denote the counterbalancing of the stimulus assignation across participants. During distractor trials, we also included the absence of audio (A_0_) or visual (V_0_) stimulation. The target stimuli consisted of either the combination of the two auditory stimuli for the audio target or the combination of the two visual stimuli for the visual target. **B**. The distractor trials consisted of an auditory period (500 ms) and a visual period (600 ms), during which either an auditory or a visual event occurred. Notably, when it was delivered as audio or visual event the absence of stimulation, we still maintained their designated durations. After the visual period, an inter-trial interval (ITI) of 3 ± 0.5 seconds occurred, and then the next trial started. The target trials consisted of the presentation of only one of the two target stimuli, followed by the ITI. **C**. Transition matrices illustrating the probability structure underlying the associations between audio-visual events in the three sessions, namely Habituation, Acquisition and Extinction. **D**. Graphical illustration of the distribution of the experimental sessions across the 6 days in which the experiment was performed. Electroencephalography (EEG) was recorded in the first and last day. **E**. Event-related potentials (ERPs) of the audio and visual events, color-coded for the experimental sessions. The audio ERPs are averaged across channels within a temporal region of interest (ROI), while the visual ERPs across an occipital ROI, both showed in the channel layout plot in the bottom-right part of the plot in each row. Shaded areas indicate SEM across participants. Dotted lines indicate the onset of the audio and visual period, denoted by the ear and eye symbol, respectively. The time in the x-axis of the visual events is referred to the onset of the audio period. On top of each lineplot, there are topoplots showing the scalp distribution, averaged across sessions, of the neural activity averaged in time windows of 250 ms. **F**. Lineplots showing the Global Mean Field (GMF) for all nine joint audio-visual events. Black shaded areas indicate SEM across participants. Green and pink highlighted temporal regions indicate the onset and offset of the audio and visual period, respectively.

Each distractor trial comprised two distinct periods, an auditory period and a visual period, during which either an auditory or a visual event was presented (Fig. 1B). In other words, there were a total of 9 joint audio-visual events that could happen in the distractor trials. Conversely, target trials consisted of either the presentation of the audio target or the visual one. Crucially, the ability to predict the visual events from the auditory cues in the distractor trials had no influence on task performance, as the target stimuli occurred in separate target trials, and their timing resembled that of the distractor audio events, offering no predictive advantage. In other words, once participants recognized that a trial did not contain a target, any predictions about the forthcoming sensory event held no task relevance.

The experiment was structured into 3 sessions, namely Habituation, Acquisition and Extinction (Fig. 1C). In the Habituation and Extinction sessions, the transition probability matrix from the audio events to the visual events was uniform, while in the Acquisition session there was a clear association between audio-visual events. These audio-visual sensory associations were systematically manipulated to probe different patterns of audio cue and visual target relationships. Specifically, we designed the transition structure to include three distinct cases. In one case, the absence of a distractor cue (A_0_) predicted the presence of a visual stimulus (V_2_). In another case, the presence of a cue (A_2_) predicted the absence of a visual stimulus (V_0_), while in the last case there was a canonical association between the presence of a cue (A_1_) and the presence of a visual stimulus (V_1_). The whole experiment was conducted over 6 days within a 2-week period, where we recorded the EEG on the first and last day (Fig. 1D).

We started the data analysis by inspecting the EEG data, plotting the average evoked potentials of the six audio-visual events (Fig. 1E) across the experimental sessions. We observed the time courses of the neural activity in the temporal and occipital regions of interest (ROI) for audio and visual events, respectively, being very similar across sessions. The topoplot averaged across sessions indicated a prominent parietal activity in the audio cue period and occipital activity in the visual outcome period. Then, we plotted the global mean field (GMF) for all audio-visual joint events to inspect the general aspects of the EEG data averaged across all sessions (Fig. 1F). The trends matched our expectations, with less GMF whenever there was the absence of sensory stimulation compared to when the stimuli were delivered.

### Decoding incidental learning effects in the neural activity

Next, we used multivariate decoding analyses (*92*) to investigate the establishment of incidental associative learning at the neural level. In other words, we operationalized whether learning of these incidental sensory associations occurred by performing classification analysis between the Habituation and Extinction sessions (i.e., before and after learning) among the audio and visual events. In the audio period (Fig. 2A), we found a robust decoding accuracy for A_1_ and A_2_ cues across the whole period (both P < 0.003 cluster-corrected, peak unbiased Cohen’s d > 4.41), peaking both around 200-300 ms post audio period onset. Searchlight analyses highlighted a significant widespread contribution of channels in decoding these learning effects. No significant decoding accuracy was observed in the absence of audio cue stimulation A_0_.

**Fig. 2.**
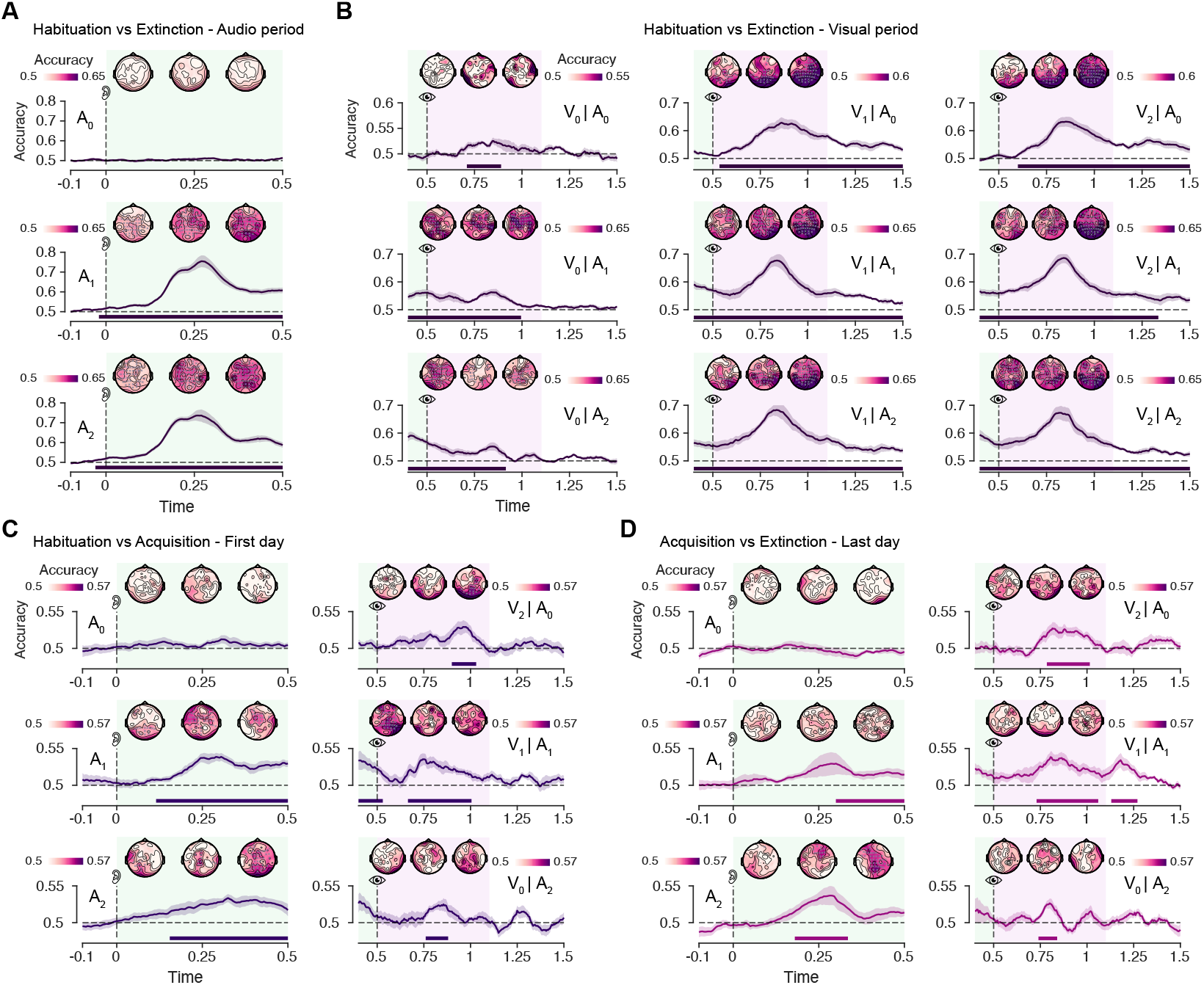
Multivariate decoding between experimental sessions across learning. Decoding accuracy between Habituation versus Extinction in the audio period (**A**) for each audio event and in the visual period (**B**) for all nine visual events conditioned on the audio events. In **C**. and **D**., decoding accuracy between Habituation versus the Acquisition session from the first day and Acquisition from the last day versus Extinction, respectively. Both results are shown in the audio period for each audio event and visual period only for the visual events having enough trials (i.e., the ones associated at 90% in the Acquisition session). Shaded areas in the lineplots indicate SEM across participants. Dotted lines indicate the onset of the audio and visual period, denoted by the ear and eye symbol, respectively. The time in the x-axis of the visual events is referred to the onset of the audio period. Green and pink highlighted temporal regions indicate the onset and offset of the audio and visual period, respectively. On top of each lineplot, there are topoplots showing the scalp distribution of the decoding accuracy averaged in time windows of 150 ms. Statistically significant channels are denoted with a grey cross in the topoplots, while horizontal bars denote statistical significance in the lineplots.

In the visual period (Fig. 2B), we performed the classification separately for each of the visual events conditioned on the audio cue events (i.e., all nine audio-visual joint events). Crucially, we found significant decoding accuracy in all these cases. Specifically, decoding accuracy was significantly above chance level across the whole visual period in all the audio-visual events where there was the presence of the visual stimulus V_1_ and V_2_ (V_1_|A_0_, V_2_|A_0_ and so on, all P < 0.003 cluster-corrected, d > 2.77). Searchlight analyses revealed how these learning effects could be decoded mostly across occipital channels. Notably, we could decode the presence of a learning effect also in absence of visual stimulation (i.e., V_0_|A_1_ and V_0_|A_2_, both P < 0.003 cluster-corrected, d > 2.37), even in the case of complete “sensory silence” where no audio or visual stimulation were delivered (V_0_|A_0_, P = 0.004 cluster-corrected, d = 2.28).

These results suggest that participants acquired the audio-visual associations, despite reporting no awareness of the learning process. However, the classification between the Habituation and Extinction sessions could be influenced by confounding factors related to the fixed temporal order of the sessions, such as fatigue, attentional drift, or general time-related effects, rather than learning per se. To control for this, we performed an additional set of classification analyses: one comparing Habituation and Acquisition sessions on the first day, and another comparing Acquisition and Extinction sessions on the last day. This approach allowed us to disentangle neural signatures of learning from those potentially attributable to the mere passage of time. Crucially, the order of learning phases was reversed across these comparisons, Acquisition followed Habituation in the first case, but preceded Extinction in the second, enabling a more rigorous assessment of learning-related neural changes independent of session chronology. Given the limited trial availability for some audio-visual pairings in the Acquisition, in the visual period we only performed the classification using the high probable audio-visual events (i.e., V_2_|A_0_, V_1_|A_1_, and V_0_|A_2_).

In the first-day analysis (Fig. 2C), we observed significant decoding accuracy in the auditory period for A_1_ (P < 0.003 cluster corrected, d = 2.14) and A_2_ (P < 0.003 cluster corrected, d = 1.84) cues, with accuracy rising well above chance following the onset of the auditory stimulus. These effects were distributed across widespread scalp regions, as revealed by the corresponding searchlight topographies. No significant decoding was found for the absence of an auditory cue (A_0_, P = 0.262 cluster corrected), consistent with the previous analysis. In the visual period, we decoded learning effects in all pairing cases (V_2_|A_0_, P = 0.035, d = 1.14; V_1_|A_1_, P = 0.004, d = 1.41; V_0_|A_2_, P = 0.031, d = 1.48; all P cluster corrected), around 250-500 ms after visual period onset and predominantly from occipital channels.

In the last-day analysis (Fig. 2D), a similar pattern emerged. Significant decoding accuracy was observed for A_1_ (P = 0.012 cluster corrected, d = 1.06) and A_2_ (P = 0.012 cluster corrected, d = 1.40) cues in the auditory period, with no reliable decoding for A_0_ trials (P > 0.05 cluster corrected). In the visual period, classification was again successful in all pairing cases with similar latencies, including both cases where a visual stimulus was present (V_2_|A_0_, P = 0.004, d = 1.26; V_1_|A_1_, P = 0.008, d = 1.53; both P cluster corrected) and absent (V_0_|A_2_, P = 0.039 cluster corrected, d = 1.72). Together, these results replicated the decoding signatures of incidental associative learning across both temporal directions, showing that the observed neural differences indeed reflected learning-related changes rather than session-related confounds.

### Fitting ideal observer models to neural data

After establishing the presence of incidental associative learning at the neural level (*82, 85*), we next deepened more on the neural signatures of the predictive mechanisms that underlied the acquisition of these incidental sensory associations. To achieve this, we adopted a computational modelling approach to examine how the human brain encoded the predictions and prediction errors generated from the statistical structure of the audio-visual incidental associations. Specifically, we used an ideal observer model (Fig. 3A), a perceptron neural network resembling the Rescorla-Wagner model for categorical data (*12, 93, 94, 56*). This model simulated the trial-by-trial predictions and prediction errors by attempting to infer the upcoming visual event from the preceding auditory event, generating prediction and error trajectories over the trial sequence. To better characterize the underlying predictive mechanisms, we first performed a model comparison between alternative formulations of the ideal observer model.

**Fig. 3.**
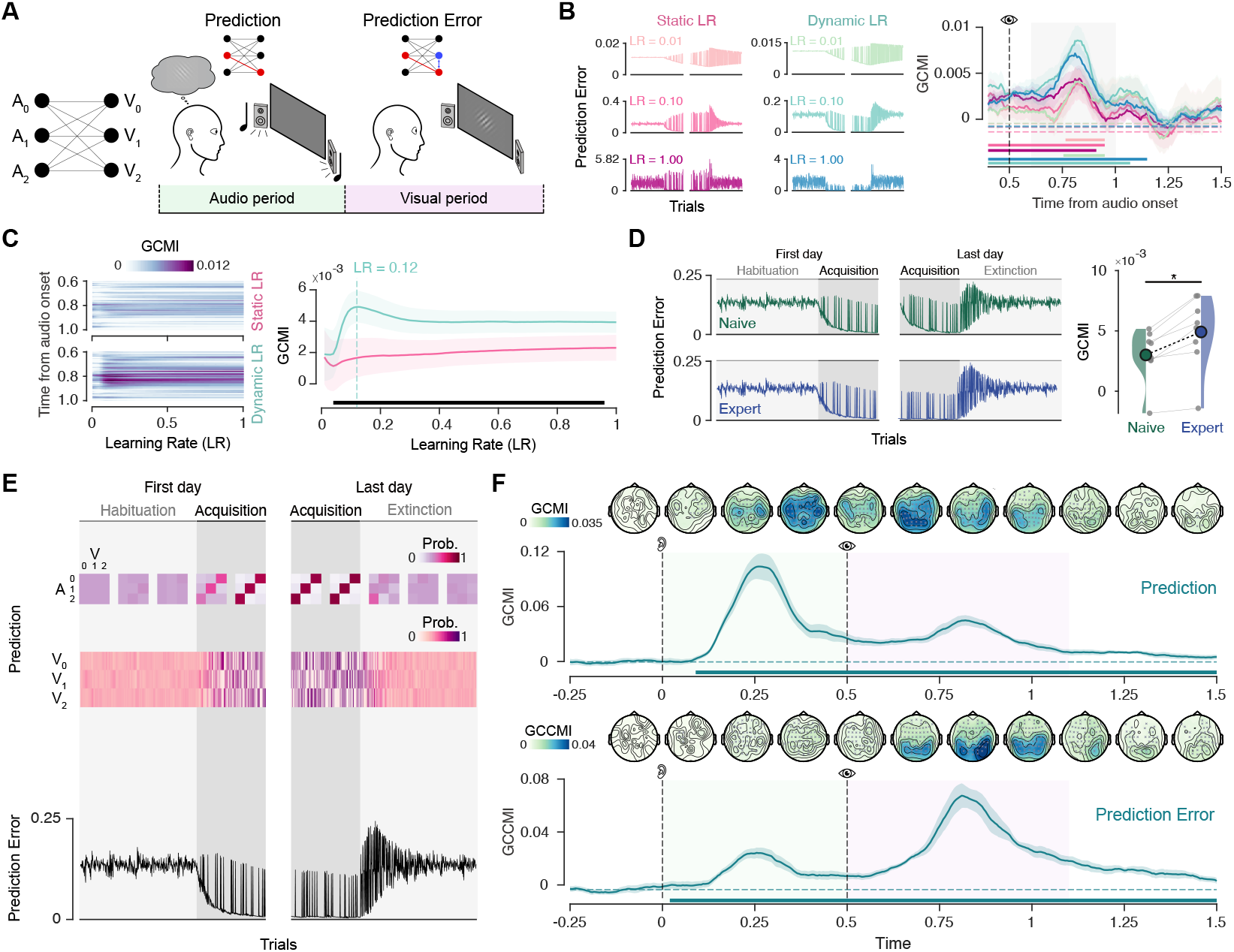
Fitting prediction and error trajectories from the ideal observer model to the neural data. **A**. Graphical illustration of the ideal observer model. On the left, schematic representation of the input and output layers of the model, while on the right an illustration of the prediction and prediction errors extracted from the model corresponding to assumed computations happening in the audio and visual periods, respectively. **B**. Model comparison between ideal observer models with static and dynamic learning rate (LR). On the left, prediction error trajectories across the first and last day for examples of model instances (example from one participant, with LR set to 0.01, 0.1 and 1). On the right, Gaussian Copula Mutual Information (GCMI) values obtained by fitting the error trajectory examples to the neural data across time in the visual period. Shaded areas represent standard error of the mean (SEM) across participants, while horizontal bars indicate statistical significance. **C**. Left, group-average GCMI values for the Static and Dynamic LR models across the time window in the visual period (from 0.6 to 1 seconds from audio period onset) and the learning rate grid search parameter, ranging from 0.01 to 1. Right, lineplots showing GCMI averaged across the selected time window as a function of the learning rate. Shaded areas represent SEM, while horizontal bars indicate statistical significance. **D**. Model comparison between naive and expert ideal observer models. Naive models are defined as those which reset their learned associations between audio-visual events after each day, while expert models keep track of those associations across the whole experiment. On the left, error trajectories for both model types for all sessions (Habituation-Acquisition of the first day and Acquisition-Extinction of the last day, example from one participant with LR = 0.12). On the right, raincloud plots showing the GCMI values (averaged in the time window of the visual period and with the best learning rate LR = 0.12) for naive and expert model. Circles indicate the mean of the distribution and dots individual participants. Asterisks represent statistical significance. **E**. Prediction and prediction error trajectories (example from one participant) extracted from the winning model (dynamic LR and expert). Top, weight matrices across experimental sessions, normalized and expressed as probabilities. Below, predictions across sessions for each visual outcome, expressed as probabilities. Bottom, prediction error trajectory across sessions. **F**. Lineplots showing GCMI values for fitting model predictions to the brain data (top) and conditional GCMI (GCCMI, conditioned on predictions) for fitting prediction errors (bottom). Shaded areas in the lineplots indicate SEM across participants. Dotted lines indicate the onset of the audio and visual period, denoted by the ear and eye symbol, respectively. Green and pink highlighted temporal regions indicate the onset and offset of the audio and visual period, respectively. On top of each lineplot, there are topoplots showing the scalp distribution of the GCMI and GCCMI values averaged in time windows of 150 ms. Statistically significant channels are denoted with a grey cross in the topoplots, while horizontal bars denote statistical significance in the lineplots.

We started by testing whether neural responses during the visual period were better captured by an ideal observer model with a static learning rate (LR), in which prediction errors are weighted uniformly over time, or by a dynamic learning rate model, where learning is adaptively modulated by the model’s epistemic uncertainty (Fig. 3B). This comparison was motivated by the Bayesian brain hypothesis (*28, 30, 87*), which posits that the brain updates its internal models of the world by integrating new sensory evidence in a statistically optimal manner. A key tenet of this framework is that prediction errors are not treated equally across time but are precision-weighted, that is, scaled according to the reliability or uncertainty of current beliefs. By fitting the prediction error trajectories to the neural data using Gaussian Copula Mutual Information (GCMI), employed here as a multivariate measure of model fit (*56, 95*), we found that the dynamic LR model consistently outperformed the static LR model (Fig. 3C), yielding significantly higher goodness-of-fit across nearly the entire learning rate parameter space (P < 0.004 cluster corrected, d = 1.69). Crucially, the winning dynamic LR model exhibited a clear peak in GCMI values at a learning rate of 0.12, allowing us to identify it as the best-fitting parameter and use it for subsequent analyses. Importantly, the superiority of the dynamic LR model was confirmed at the individual participant level, confirming the robustness of the result (Fig. S1A).

Next, we examined whether participants retained the acquired sensory associations across days, or whether these incidental associations were instead relearned anew on each experimental day. This analysis aimed to address whether statistical regularities, despite being task-irrelevant and implicitly learned, are nonetheless consolidated into memory and persist over extended timescales, such as across days or weeks. To this aim, we fitted neural responses in the visual period using the prediction error trajectories extracted from an “expert” ideal observer model, which retained knowledge of the learned audio-visual associations from previous days, or by a “naive” model that reset its associative memory at the beginning of each day. These models produced distinct prediction error trajectories, primarily differing in their initial error estimates on the last experimental day (Fig. 4D, left). In other words, the prediction error in the naive model was generally higher compared to the expert model during the Acquisition session of the last day. Crucially, we found that the expert model outperformed the naive model, testing with the previous best-fitting LR parameter (Fig. 4D, right, t(7) = 5.03, P = 0.001, d = 1.58) or along the parameter space (Fig. S1B, P = 0.039 cluster corrected). Notably, this trend was also confirmed at the individual participant level (Fig. S1C). These findings indicate that the brain retains in memory sensory associations, regardless of their current behavioral relevance, over extended timescales.

**Fig. 4.**
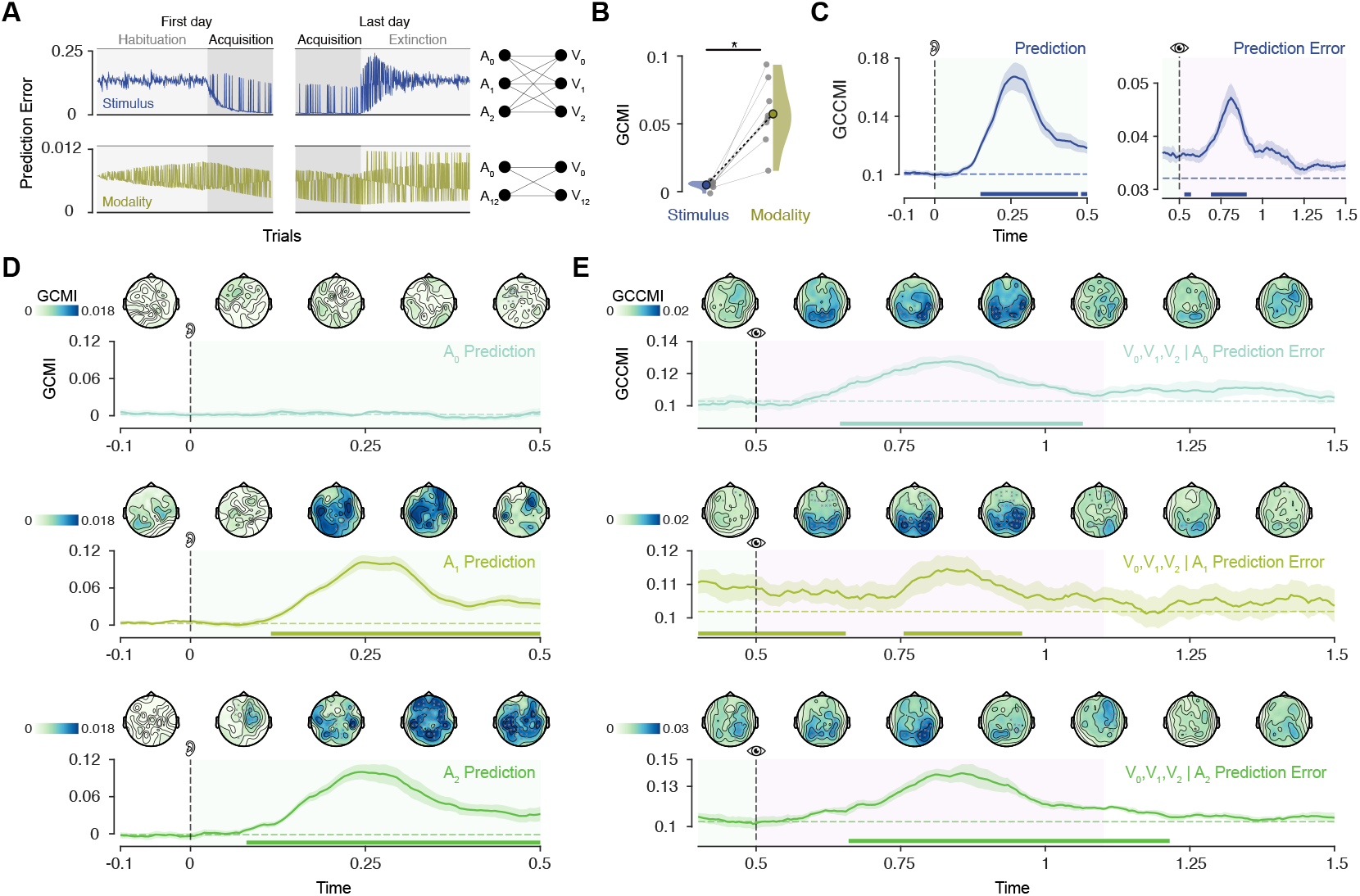
Event-specific spatiotemporal characteristics of prediction and error encoding. **A**. Model comparison between stimulus- and modality-level ideal observer models. On the left, error trajectories for both model types for all sessions (Habituation-Acquisition of the first day and Acquisition-Extinction of the last day, example from one participant). On the right, schematic representation of the input and output layers of both model types. **B**. Raincloud plots showing the GCMI values (averaged in the time window of the visual period and with the best learning rate) for stimulus and modality model. Circles indicate the mean of the distribution and dots individual participants. Asterisks represent statistical significance. **C**. Conditional GCMI (GCCMI) values by fitting prediction and error trajectories of the stimulus-level model conditioned on the modality-level model’s trajectories. Lineplots showing GCMI values for fitting the (stimulus-level model as in Fig. 3F) model predictions to the neural data (**D**) and conditional GCMI (GCCMI, conditioned on predictions) for fitting model prediction errors (**E**), specifically for each auditory cue. On top of each lineplot, topoplots show the scalp distribution of the GCMI and GCCMI values averaged in time windows of 150 ms. Statistically significant channels are denoted with a grey cross in each topoplot. For lineplots in **C**., **D**. and **E**., shaded areas represent SEM across participants. Dotted lines indicate the onset of the audio and visual period and green and pink highlighted temporal regions represent the audio and visual period, respectively. Horizontal bars denote statistical significance.

### Neural encoding of incidental predictions and prediction errors

The above results revealed that the ideal observer model that best captured participants’ neural responses was the one that incorporated epistemic uncertainty to modulate prediction errors and retained incidental sensory associations in memory across days. Having established the best-fitting model, we next asked: how are these incidental predictions and prediction errors encoded in the brain? To address this question, we extracted the trial-by-trial predictions and error signals from the winning model (Fig. 3E) and fitted them to the neural data using a time-resolved and searchlight-based approach. Crucially, we found significant results when we tried to fit neural responses with both prediction and error trajectories (Fig. 3F).

Incidental predictions were encoded approximately 250 ms after audio period onset (P < 0.003 cluster corrected, d = 3.64), primarily distributed over fronto-central channels. In contrast, incidental prediction errors were encoded around 300 ms after visual period onset (P < 0.003 cluster corrected, d = 3.22), and were strongly localized over occipital sensors. Notably, prediction errors were fitted to the neural data using conditional GCMI (GCCMI), which quantifies the unique contribution of prediction errors after accounting for predictions. This step was essential because predictions and prediction errors are often correlated, due to both the statistical structure of the sensory input and the computations implemented by the ideal observer model. This correlation is also reflected in the temporal overlap of prediction- and error-related encoding, with residual signal observed in the visual period for predictions and in the auditory period for errors. Importantly, because prediction errors cannot be computed prior to the onset of the visual input, early encoding of error signals likely reflects this underlying correlation, further motivating the use of a conditional approach.

For completeness, we also fitted the neural data with the prediction error trajectories using GCMI, without conditioning on predictions (Fig. S1D). This analysis revealed a broadly similar temporal profile to that observed with the conditional approach, with prediction error signals emerging around 300 ms after visual period onset (P < 0.003 cluster corrected, d = 1.54 peaking post visual period onset) and being distributed across posterior scalp regions. However, compared to the GCCMI results, the use of standard GCMI resulted in more widespread and temporally diffuse effects, particularly in the early audio period where actual prediction errors could not yet be computed. In sum, we characterized the temporal and spatial neural signatures underlying the encoding of incidental predictions and prediction errors that support the acquisition of task-irrelevant sensory associations in the human brain.

### Stimulus-specific versus sensory modality predictions and errors

After this crucial step, we tested whether the observed neural encoding of predictions and prediction errors reflected truly stimulus-specific (or event-specific) associations or, instead, more general associations at the level of sensory modality. We compared the original stimulus-specific ideal observer model with a reduced model that tracked only the presence or absence of sensory stimulation (Fig. 4A). We first evaluated whether the modality-level model was best fit by a static or dynamic learning rate, using the same model fitting procedure applied to the stimulus-level model. Both model variants showed similar overall support from the data, with goodness-of-fit peaking at a learning rate of 0.01. However, the static model outperformed the dynamic model at higher learning rates (P < 0.003 cluster-corrected) and was more consistently favored at the individual participant level (Fig. S2A–C). Thus, we selected the static LR version for the modality-level model for subsequent analyses.

We then directly compared the predictive power of the stimulus-specific and modality-level models. Fitting both models’ error trajectories to the neural data during the visual period revealed significantly higher GCMI values for the modality-level model compared to the stimulus-specific model (Fig. 4B, t(7) = 5.78, P < 0.001, d = 1.81). This finding was in line with our expectations, as predicting the sensory modality is computationally simpler and more general than predicting specific stimuli. Critically, however, what we were primarily interested in was whether the modality-level model could fully account for the variance in neural responses, or whether there remained residual variance specifically explained by the stimulus-specific model.

To test this, we used conditional mutual information (GCCMI) to fit the predictions and errors from the stimulus-specific model while controlling for the corresponding trajectories from the modality-level model. This analysis revealed that, even after accounting for the information captured by the modality-level model, the stimulus-specific model continued to explain a significant portion of the neural data (Fig. 4C), both for predictions (P = 0.004 cluster corrected, d = 2.64) and prediction errors (P < 0.003 cluster corrected, d = 2.67). These results were confirmed even when we equated the learning rate parameter of the modality-level model to the stimulus-specific model (Fig. S2D, resetting the LR to 0.12), with both stimulus-specific predictions (P = 0.008 cluster corrected, d = 2.67) and prediction errors (P < 0.003 cluster corrected, d = 2.45) still accounting for a significant portion of the residual variance after controlling for modality-level variables (Fig. S2E). In sum, these findings demonstrate that the neural encoding of predictions and prediction errors was not limited to broad modality-level expectations, but also reflected fine-grained, stimulus-specific associations that participants acquired incidentally over the course of learning.

### Event-specific neural signatures of prediction and error encoding

Finally, after establishing the overall encoding of incidental predictions and prediction errors, we examined the neural encoding separately for each auditory event cue. Given that each audio cue in our design was associated with a distinct pattern of sensory association (A_0_→V_2_, A_1_→V_1_, and A_2_→V_0_), we aimed to determine whether predictions and errors were consistently encoded across all these cases. This more granular analysis allowed us to verify whether the encoding observed at the overall level generalized to the specific types of audio-visual associations embedded in the task, which could not be distinguished in the previous, aggregated analyses.

By fitting the trial-by-trial prediction and error trajectories extracted from the winning ideal observer model (stimulus-specific, expert and dynamic LR) separately for each auditory cue to the neural data, we found that incidental predictions (Fig. 4D) were robustly encoded around 250 ms after audio period onset for both A_1_ (P < 0.003 cluster corrected, d = 2.87) and A_2_ (P < 0.003 cluster corrected, d = 3.30) cues, with maximal GCMI values over fronto-central channels. No significant prediction encoding was found following the absence of an auditory cue (A_0_, P > 0.05 cluster corrected), consistent with the decoding results.

In contrast, incidental prediction errors (Fig. 4E), fitted to the neural data after accounting for predictions using GCCMI, were prominently encoded around 300 ms after the onset of the visual period, predominantly over occipital channels, across all three sensory association types: prediction of a visual stimulus following no auditory cue (A_0_→V_2_, P < 0.003 cluster corrected, d = 1.80), prediction of a visual stimulus following an auditory cue (A_1_→V_1_, P = 0.016 cluster corrected, d = 1.19), and prediction of no visual stimulus following an auditory cue (A_2_→V_0_, P < 0.003 cluster corrected, d = 1.93). These patterns were further confirmed when using standard GCMI without conditioning on predictions, showing similar latency effects and spatial distribution on the scalp in the searchlight results (Fig. S3, all P < 0.004, d > 1.77). In sum, these findings reveal that the brain dynamically tracks fine-grained sensory associations, encoding both predictions and prediction errors across different event structures, involving the absence of cues or expected sensory input.

## Discussion

Our results provide compelling evidence that the human brain acquires sensory associations independently of their immediate behavioral relevance, while also revealing the predictive coding mechanisms that underpin the acquisition of these incidental associations at the neural level. Our findings accord well with previous studies demonstrating that task relevance is not strictly required for the brain to build predictive models of the environmental regularities (*80, 82, 84, 85*). Importantly, our results extend previous work in several directions by showing that both predictions and prediction errors were robustly encoded in neural activity and characterizing their distinct spatiotemporal neural profiles. For instance, in our study the sensory associations remained consistently task-irrelevant, unlike in previous studies (*83, 84*) where predictive cues were relevant in some blocks and irrelevant in others. Such alternation may inadvertently encourage participants to retain associations across tasks, introducing potential covert task relevance.

Interestingly, while previous work by den Ouden et al. (*82*) demonstrated prediction error signals underlying incidental sensory associations, they found no evidence for incidental associative learning when the absence of a cue predicted the presence of a visual stimulus. In contrast, our results show that participants acquired incidental sensory associations in which the absence of an auditory cue predicted the occurrence of a visual event, as well as associations in which the presence of a cue predicted the absence of subsequent visual stimulation (*85*). Notably, we found that the latency and spatial distribution of prediction and prediction error encoding were remarkably consistent across different types of sensory associations, regardless of whether they involved the presence or absence of sensory stimulation. Across both multivariate decoding and computational modeling results, we did not find evidence for neural encoding of incidental sensory predictions when the cue was the absence of auditory stimulation. This may reflect the fact that, without an external sensory input, it is difficult with no discrete event to time-lock predictive processes to, particularly in our paradigm, where stimuli were embedded in a continuous stream without explicit trial onsets. Alternatively, the absence of input may not evoke a sufficiently specific sensory representation to be detectable with EEG. Crucially, however, we did observe robust encoding of prediction errors in these trials, likely because a sensory event, a visual stimulus in this case, was subsequently (mostly) delivered.

Building on the idea that the predictive mechanisms optimizing stimulus–response learning may also support perceptual learning of stimulus–stimulus associations (*27, 29, 82*), our findings extend this framework by showing that predictive learning can generalize beyond physical contingencies to encompass general sensory events, including the absence of stimulation itself. This challenges a common assumption in associative learning theories, that the physical presence of stimuli is necessary for associations to form (*90*). Instead, our results align with theoretical proposals emphasizing that informativeness, rather than simple temporal contiguity or physical presence, governs the formation of predictive associations in the brain (*6, 89, 91, 96*). This view was already anticipated in foundational work by Rescorla (*11, 88, 89*), who demonstrated that temporal pairing between stimuli and outcomes is neither necessary nor sufficient for associative learning. Through carefully controlled experiments involving inhibitory classical conditioning paradigm, he showed that it is the contingency, the degree to which one event provides information about the likelihood of another, that determines whether sensory associations form. When the presence or absence of a cue systematically predicts an outcome, as in our experimental paradigm, even in the absence of strict temporal contiguity, associative learning can occur.

Beyond demonstrating the incidental acquisition of sensory associations, our results provide novel insights into the computational principles governing how these associations are learned. Specifically, by modeling trial-by-trial neural 7responses with ideal observer models, we found that prediction errors were not weighted uniformly over time. Instead, learning was best captured by a dynamic learning rate model, in which prediction errors were scaled according to the model’s epistemic uncertainty. This finding aligns with the Bayesian brain hypothesis (*28, 30, 87*), which posits that the brain updates its internal models in a statistically optimal manner by adjusting learning rates based on the reliability of prior beliefs. By showing that epistemic uncertainty modulates prediction error processing even during the incidental acquisition of task-irrelevant sensory associations, our results extend Bayesian accounts of learning beyond explicitly goal-directed contexts.

In addition to dynamic updating, our findings demonstrate that incidental sensory associations are not transient but are retained in memory across days. Our findings align with recent behavioral evidence demonstrating that incidentally acquired information can persist in long-term memory even when it is unattended and task-irrelevant (*97*–*100*). For example, Hutmacher and Kuhbandner (*99*) showed that auditory stimuli presented outside the focus of attention, and irrelevant to task demands, were nonetheless encoded in sufficient detail to support recognition after a 24-hour delay. Similarly, Gabay et al. (*100*) found that incidental exposure to novel acoustic categories during an unrelated visuomotor task led to robust category learning that became evident only after offline consolidation. These findings support the existence of a perceptual memory system capable of storing detailed sensory information independently of top-down goals, attention, or awareness. Our results extend this literature by showing the neural mechanisms underlying incidental learning and for cross-modal sensory associations, showing that not only is incidental information consolidated over days, but that it continues to shape neural representations in a way consistent with predictive coding theories (*3, 33, 36, 56, 85*). Together, these behavioral and neural findings suggest that the brain preserves incidental sensory regularities in long-term memory systems, providing a latent store of statistical knowledge that could be leveraged if environmental contingencies later become behaviorally relevant.

In conclusion, our results provide empirical support for a longstanding question in neuroscience, whether the human brain acquires sensory associations through incidental learning. By demonstrating that such associations are learned, retained across days, and integrated into predictive neural computations, our findings point to a broader principle, namely, that the brain seeks to reduce uncertainty about environmental state transitions, regardless of immediate behavioral demands. This suggests that constructing predictive models is not merely a means to achieve specific goals, but a core computational imperative of brain function. Understanding this intrinsic drive may be key to explaining how the brain builds flexible internal models of the world that support perception, learning and adaptive behavior.

## Methods

### Participants

Eight volunteers (4 females, mean age 26.7, range 22-38) participated in this study. All were right-handed with normal or corrected-to-normal vision and normal hearing, had no history of neurological disorders and were not taking any neurological medications. All participants gave informed written consent. The study was conducted in accordance with the Declaration of Helsinki and approved by the University of Trento Ethics Committee (protocol number 2018-009).

### Procedure

Participants were instructed to perform a target detection task by being exposed to a stream of audio and visual stimuli while sitting in a dimly lit booth at a distance of 1 m from the CRT monitor (22.5 inch VIEWPixx; resolution: 1024 × 768 pixels; refresh rate: 100 Hz; screen width: 50 cm). The task consisted of pressing a button upon detecting a target stimulus, while ignoring the distractor stimuli (Fig. 1A). The experimental script was generated using OpenSesame with PsychoPy as backend (*101, 102*).

Each distractor trial consisted of two distinct periods (Fig. 1B): an auditory period (500 ms) and a visual period (600 ms), during which either an auditory or a visual event occurred. Importantly, these periods could also include the absence of stimulation while still maintaining their designated durations. During the auditory period, one of three possible events occurred (Fig. 1A). Two were auditory stimuli (A_1_ and A_2_), low and high frequency tones of 375 Hz and 750 Hz (roughly corresponding to a fourth and fifth octave F_#_ musical note), and the third was the absence of auditory stimulation, denoted as A_0_. Similarly in the visual period, one of three possible events occurred (Fig. 1A). Two were visual stimuli (V_1_ and V_2_), Gabor patches (4.4° × 3.4° visual angle) with Gaussian envelope, standard deviation of 18.0 and a spatial frequency of 0.08 cycles/pixel displayed in a grey background (RGB: [128, 128, 128]), one with 45° orientation (right) and the other one with 135° orientation (left). The third visual event was the absence of visual stimulation, denoted as V_0_. The assignment of the stimuli to the audio and visual events (besides the absence of stimulation) was counterbalanced across the participants. After the visual period, an inter-trial interval (ITI) of 3 ± 0.5 seconds occurred, and then the next trial started. On target trials (Fig. 1B), there was presented only one of the two target stimuli, consisting of either the combination of the two auditory stimuli for the audio target or the combination of the two visual stimuli for the visual target (Fig. 1A), followed by the ITI. The trials were grouped into blocks of 102 trials, with 12 target trials (6 for each sensory modality) randomly interspersed between the remaining 90 distractor trials. Between two consecutive blocks, a short pause of about one minute was delivered.

Unbeknownst to them, we manipulated the statistical associations between the distractor stimuli. The experiment was structured into 3 sessions, namely Habituation, Acquisition and Extinction, conducted over 6 days within a 2-week period. In other words, while the six experimental days were not evenly spaced for all participants, every participant completed the experiment within this two-week timeframe. In first day, participants were exposed to the Habituation session (5 blocks) and the start of the Acquisition session (3 blocks), which continued over the course the following four days (8 blocks each). In the last day, participants were exposed to the end of the Acquisition session (3 blocks) and the Extinction session (5 blocks). EEG recordings were collected during the first and last day (Fig. 1D).

In the Habituation and Extinction sessions (Fig. 1C), the transition probability matrix from the audio events to the visual events was uniform (i.e., all nine joint events were equally probable).

Conversely, in the Acquisition session the absence of audio stimulation (A_0_) predicted the presence of the V_2_ stimulus with a 90% probability and V_1_ with 10% probability. Moreover, A_1_ predicted strongly V_1_ (90%) and weakly the absence of visual stimulation (V_0_, 10%) and A_2_ predicted V_0_ with a 90% probability and V_2_ with 10% probability. All the remaining joint events were not occurring during the Acquisition session. Importantly, the predictive relationships between the distractor stimuli were completely irrelevant for detecting the target stimuli. To assess whether participants had noticed these statistical associations, they were debriefed at the end of the experiment using a questionnaire. The questionnaire included two questions. The first asked whether they had noticed anything about the stimuli, while the second explicitly inquired whether they had perceived any patterns among the distractor stimuli. Crucially, participants did not report awareness of these associations.

### EEG data acquisition and pre-processing

EEG data were recorded from a standard 10-5 system with 64 Ag/AgCl electrodes cap (EasyCap, Brain Products, Germany) at a sampling rate of 1 kHz. Impedance was kept below 10 kΩ for all channels. AFz was used as the ground and the right mastoid was used as reference. We first segmented the data into epochs from -2 to 3.5 seconds relative to the onset of the audio period in the distractor trials, to account for edge artifacts due to subsequent filtering operations. Then, we applied a fourth-order Butterworth high-pass filter at 1 Hz and a notch band-stop filter at 49-51 Hz to remove line noise. Next, data were resampled to 200 Hz and re-referenced to a common median reference. We applied Independent Component Analysis (ICA) to decompose the signal and discard stereotypical artifacts (*103*) such as eye movement, muscular and cardiac artifacts, using FastICA (*104*). The data we input for the ICA decomposition were copied and high-passed at 2 Hz to improve the decomposition by reducing slow drifts and enhancing the stationarity of the data. Then, the unmixing matrix was applied to the prior data and artifactual independent components were pruned. Finally, the data were re-epoched from -0.5 to 1.5 seconds relative to the onset of the audio period and baseline correction was applied between −0.5 and 0 seconds. As a sanity check, we computed the evoked response potentials (ERP) of the audio and visual events across all sessions and plotted them as lineplots in the temporal and occipital region of interests (ROI) and as topoplots across time in steps of 250 ms (Fig. 1E). We also plotted the global mean field (GMF) of the data across all sessions for all the nine joint audio-visual events (Fig. 1F). Both ERP and GMF were temporally smoothed with a moving average window of 100 ms.

### Multivariate decoding

Next, we applied multivariate decoding (*92*) to classify audio and joint audio-visual events between Habituation and Extinction sessions (Fig. 2). We also classified audio events and specific joint audio-visual events conditioned on the trial numerosity (i.e., the audio-visual associations that were paired at 90% in the Acquisition session) between Habituation and Acquisition on the first day EEG recording and between Acquisition and Extinction in the last day. For each participant, we used Linear Discriminant Analysis (LDA) as classifier and a stratified 5-fold cross-validation scheme repeated 3 times to evaluate the classification performance. Accuracy was used as a metric of model performance. To avoid spurious performance given by class sample imbalances, we randomly undersampled the trials of the majority class to match the numerosity of the minority class in the training set for each fold. We ran decoding analyses across the time dimension, using all channels as features for each time-point, to obtain a time-resolved measure of decodability. Time-resolved decoding accuracy was temporally smoothed with a moving average window of 100 ms. We also performed decoding in a searchlight fashion (*105*) to investigate the spatial distribution of the discriminative effects, using as features channel neighbors in a radius of 4 cm and averaging the signal across time in specific time windows. Time windows were defined from the onset of either the audio or visual period in steps of 150 ms.

### Ideal observer model

Then, we used an ideal observer model to investigate how predictions and prediction errors are encoded in the neural activity (*56, 93, 94*). The employed ideal observer model can be conceived as a perceptron neural network (Fig. 3A), receiving as input one audio event at a time (at the trial level) and attempting to predict the next visual event. We represented the audio events categorically as a one-hot encoded vector of the same length as the number of different audio events present in the stimulation sequence *x*_*t*_ ∈ ℝ^1×*m*^, where *n* is equal to 3 and *t* indexes the trial in the sequence. We also represented the visual events as one-hot encoded vectors *y*_*t*_∈ ℝ^1×*m*^, where *m* is equal to 3 since there was three possible visual events to predict. The trainable model parameters were encoded in a weight matrix *W*_*t*_ ∈ ℝ^*n*×*m*^, which connected the input and output layer. The weight matrix was initialized as a uniform prior over the categorical distribution of the visual events, with all values having *m*^−1^ as entries. Given each audio event, the model predicted the next visual event according to the following equation:

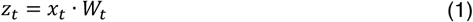

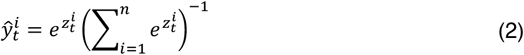

where ŷ_*t*_ represents the prediction of the model for the next visual event. We defined the loss function ℒ

as the maximum likelihood or cross-entropy function:

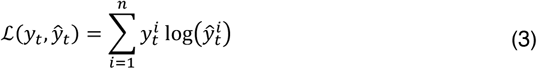

The model was trained by passing all the audio events from the stimulus sequence at a trial and after each observation, we computed the partial derivative of the loss function with respect to *W*_*t*_:

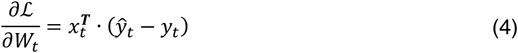

This gradient, combined to a free learning rate parameter α, gave rise to our measure of prediction error:

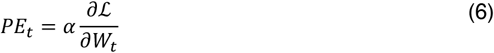

Finally, we updated the parameters *W*_*t*_ using the gradient descent algorithm:

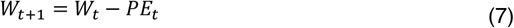

Notably, this model can also be viewed as a categorical version of the Rescorla-Wagner model (*12*). Once we sequentially passed to this ideal observer model the same audio-visual stimuli that we presented to each participant, we extracted from the model the prediction error trajectory (*56*). Thus, we fitted these error trajectories to the neural responses to test whether participants learned the statistical associations between the audio-visual events, assuming that associative learning was mediated by prediction errors as formulated by predictive coding theories of learning. We used this model to firstly test whether a static or a dynamic learning rate (LR) would better predict neural responses (Fig. 3B). We defined a static LR model the one which was parametrized by a constant α parameter throughout the whole experiment. Conversely, a dynamic LR model was parametrized by an α^*dyn*^ parameter that depends both on a fixed scaling factor α and on the current epistemic uncertainty of the model on the environmental transition dynamics, operationalized as the Shannon entropy of the predictive distribution ŷ_t_

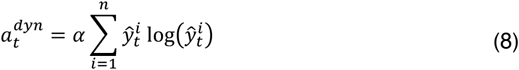

Higher entropy reflects greater uncertainty about upcoming events, leading to larger updates in belief, while lower entropy, indicating more confident predictions, results in smaller updates. This way of modeling the learning dynamics is inspired by Bayesian models of learning (*20*), in which optimal belief updating is achieved by weighting the prediction errors by the precision of their current belief. Model comparison between the static and dynamic LR models was performed with a grid search, with the α learning rate parameter ranging from 0.01 to 1 in steps of 0.01 with a total of 100 tested model instances for each model type. We fitted the prediction error trajectories coming from these model instances to the whole brain in the time window ranging from 100 ms to 300 ms after the onset of the visual period (i.e., 600 to 1000 ms after the onset of the auditory period).

As a measure of goodness of fit, we adopted the Gaussian Copula Mutual Information (GCMI) method, a robust multivariate statistical framework combining the statistical theory of copulas with the analytical solution for the Shannon entropy computation of Gaussian variables (*95*). This approach imply that each variable (brain data and prediction error trajectories) is transformed into a Gaussian variable using the inverse normal transformation. The inverse normal transformation for the neural data was applied to each feature univariately. The transformed values are obtained as the inverse standard normal cumulative distribution function (CDF) evaluated at the empirical CDF values of each variable. After this procedure, the mutual information is computed parametrically for Gaussian variables as following:

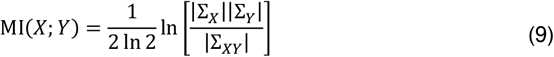

where Σ_*X*_ and Σ_*Y*_ are the covariance matrices of *X* and *Y*, respectively and Σ_*XY*_ is the covariance matrix for the joint variable (*X, Y*). Notably, this quantity was corrected for the sampling bias (*95*). Here, the *Y* variable denoted the (univariate) prediction error trajectory and *X* denoted the multivariate neural data. GCMI values were computed in a time-resolved fashion (*92*), using all channels as variables for each time point in the tested temporal window (Fig. 3C). Then, we averaged the GCMI values across all time-points and compared the two model types across the different learning rates.

Next, we also performed model comparison between an ideal observer model that stored the learned relationships across sessions or another model that it was restarting the learning process at every session. In other words, we investigated whether in the last session the neural responses could have been best described by a “naive” model that started that session with a uniform prior for predicting the visual events from the audio ones, or by an “expert” model that started the last session with already some prior knowledge about the underlying probabilistic structure of the audio-visual stimuli associations. Thus, we extracted the prediction error trajectories from the expert model, which was the one that won the previous model comparison (i.e., the dynamic LR model with a learning rate parameter of 0.12), and from the naive model, which was the same model but with the weight matrix reinitialized with a uniform prior at the start of the last session. We compared the two models by using the same model fitting procedure as above (Fig. 3D).

Then, we used the winning model to investigate the spatiotemporal characteristics of prediction and prediction error encoding in the brain underlying the acquisition of the incidental sensory associations (Fig. 3E). We used the same approach we used in the multivariate decoding analysis, by applying the GCMI to fit these model prediction and error trajectories either in a time-resolved or searchlight fashion, with the same hyperparameters as above. For predictions, we used the *y*-quantities extracted from the model for each trial, resulting in a matrix with dimensionality number of trials times the number of visual events. For prediction errors, we used the same prediction error trajectory as described above in the model comparison paragraph. Additionally, we computed the conditional mutual information (CMI) between the neural data, the error trajectory and the discrete categorical model predictions. This is crucial since the predictions and prediction errors are moderately correlated in these ideal observer models, mostly due to the statistical properties of the presented stimulus sequence. Therefore, this could confound the interpretation of prediction error encoding, which occurs during the visual period after the prediction is made, but not the prediction encoding itself, which takes place during the audio period. To compute the CMI between two continuous variables conditioned on a categorical one, we used the approach proposed by Ince and colleagues (*95*). We first computed the GCMI between neural data and error trajectories within each visual event class predicted by the model across trials. The final CMI is computed as a weighted average of the within visual event class mutual information terms, normalized by the empirical visual event class probabilities.

After this step, we also compared the model we used to investigate the encoding mechanisms of prediction and prediction errors with an ideal observer model that captured statistical associations between sensory modality events rather than stimulus-specific events (Fig. 4A). Essentially, we created another model that had as input only the presence or absence of audio stimuli and predicted the presence or absence of visual stimuli, effectively compressing the A_1_ and A_2_ into a single input unit A_12_ and similarly for the output layer the V_1_ and V_2_ into a single V_12_ unit. In other words, this model captured prediction and errors at the “modality” level, while the previous model was tracking statistical associations at the “stimulus” level. We estimated the best learning rate parameter for the modality model and compared between the static and dynamic version (Fig. S2A-C). Thus, we picked the best modality model and compared with the stimulus-specific model (Fig. 4B). Then, we investigated whether the modality model explained all the variance in the neural responses underlying predictions and prediction errors in the audio and visual period. To this aim, we computed the CMI between the neural data and either the prediction or error trajectory from the stimulus-specific model conditioned on either the prediction or error trajectory from the modality model, respectively (Fig. 4C). Here, we computed the CMI with the canonical form, using the GCMI estimator, since all the variables employed were univariate (error trajectories) or multivariate (neural data and prediction trajectories, trivariate for the stimulus-specific model and bivariate for the modality model) continuous variables. Notably, we also repeated this analysis by using prediction and error trajectories from a modality model having the same learning rate parameter as the stimulus-specific model.

Finally, we investigated the spatiotemporal characteristics of prediction and prediction error encoding specific for each auditory event. Prediction and error trajectories were extracted from the same model that we used for the analysis with all the auditory events aggregated (i.e., the stimulus-specific dynamic LR with stored associations across sessions). We fitted these variables to the neural data, separately for the trials associated with each auditory event, with the GCMI in a time-resolved or searchlight fashion, with the same hyperparameters as above. Moreover, we also computed the CMI between the neural data and the prediction error conditioned on the model predictions. Here, we used the continuous trivariate predictions instead of the discrete version as in the aggregate analysis since the number of trials for some specific visual event classes was too low and computed the CMI with the canonical form.

### Statistical analysis

We assessed the statistical significance of our findings from the model comparison across the parameter space, the decoding analyses and model fitting to the neural data in time-resolved or searchlight approach, using mass univariate cluster-based paired permutation tests (*106*). We determined contiguous clusters using t-tests across participants with an α threshold set to 0.05. For searchlight results, we used the same topological neighbor structure defined in the decoding analysis to define spatial contiguity, while for time-resolved analyses and model comparison across the learning rate dimension contiguity was naturally defined as the previous and next data point along either the temporal or parameter dimension. For classifier accuracy in the decoding analyses, we compared it to chance level using one-tail t-tests (the alternative hypothesis was that the accuracy was greater than chance level, H_1_ > 0.5). For model comparison, we used two-tail t-tests and directly compared the two model instances. For model fitting to the brain data results, we compared the goodness of fit measure (GCMI or GCCMI) to the average values in the baseline period (define as -0.5 to 0 seconds from audio period onset), by subtracting the baseline value from the observed values (for searchlight results, the subtraction was performed channel-wise) and perform a one-tail t-test (H_1_ > 0). We performed the permutation tests using all possible permutations (2^8^) and maxsum as cluster statistic. Each candidate cluster was assigned a p-value by comparing its cluster statistic to the distribution of cluster statistics of the random permutation’s strongest clusters, with an α set to 0.05 for determining statistical significance. Effect size was measured with the unbiased estimate of Cohen’s d, which is also known as Hedge’s g, and reported the peak value in the significant clusters. When comparing model instances across participants with a univariate measure, we used paired two-tail t-tests.

## Data Availability

Minimally preprocessed EEG data to reproduce all the results will be released in an openly available repository upon publication.

## Code Availability

Data analysis was carried out in MATLAB version 2022a. We used the open-source library Fieldtrip (version 20220827) available at https://github.com/fieldtrip/fieldtrip for EEG analysis. We also used the GCMI library (version 1.0) for information-theoretic analyses openly available at https://github.com/robince/gcmi.

## Acknowledgments

We thank Hanneke den Ouden and Charles Randy Gallistel for fruitful discussions about the neural and cognitive mechanisms underlying the acquisition of sensory associations via incidental associative learning. This study was supported by the BIAL foundation (Grant No. 137/22 to A.C.). L.B. is supported by Lundbeck Foundation (Talent Prize 2022), Center for Music in the Brain which is funded by the Danish National Research Foundation (project number DNRF117) and Nordic Mensa Fund.

## Author Contributions

A.G.: Conceptualization, Methodology, Investigation, Data curation, Software, Formal analysis, Visualization, Writing - original draft, Writing - Review & Editing

C.R.: Software, Formal analysis, Writing - Review and Editing

L.B.: Software, Formal analysis, Writing - Review and Editing

C.B.: Methodology, Writing - Review & Editing

A.C.: Conceptualization, Methodology, Writing - Review & Editing, Supervision, Project administration

## Supplementary Information

**Fig. S1.**
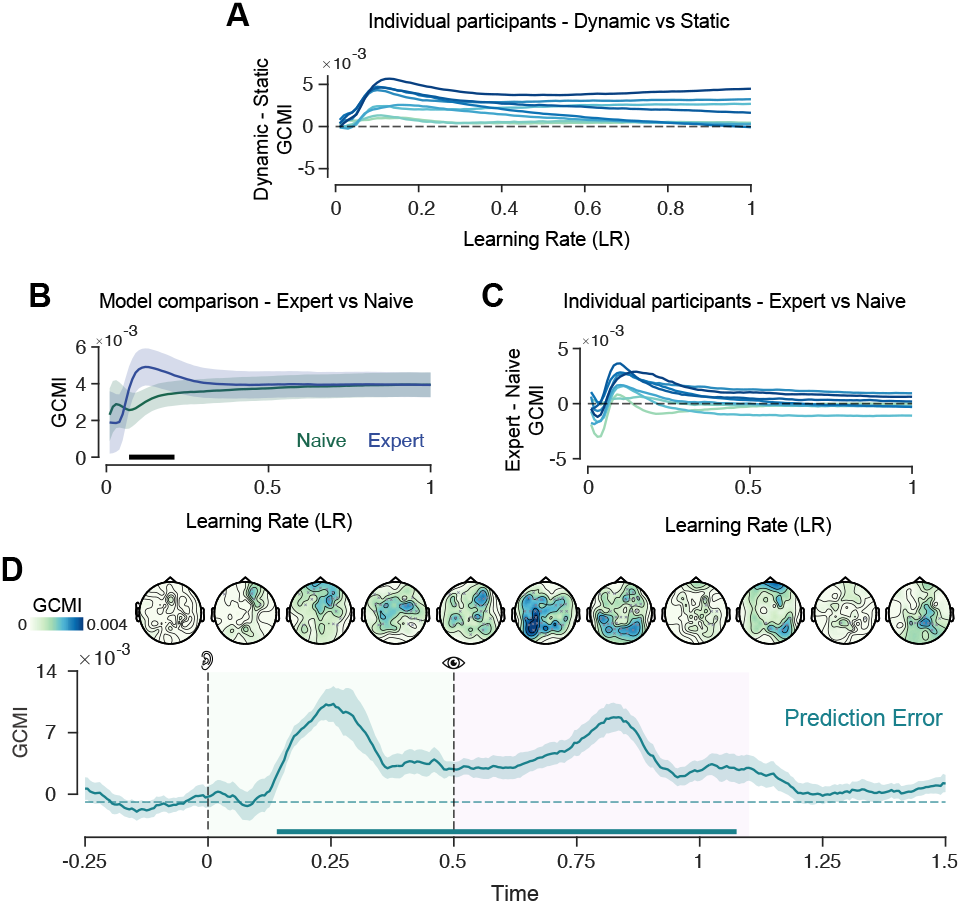
Model comparison among different ideal observer models fitted to the brain data. A. Comparison between the Static and Dynamic LR model at the participant level. Each line is a participant, showing the difference between the GCMI values of the two models as a function of the learning rate parameter. **B**. Model comparison between Naive and Expert models at the group level as a function of learning rate. Shaded areas represent SEM across participants, while horizontal bars statistical significance. **C**. Difference between the GCMI values of the Naive and Expert models, at the participant level, as a function of the learning rate. **D**. Lineplot showing GCMI values for fitting model prediction errors to the brain data. Shaded areas indicate SEM across participants. Dotted lines indicate the onset of the audio and visual period, denoted by the ear and eye symbol, respectively. Green and pink highlighted temporal regions indicate the onset and offset of the audio and visual period, respectively. On top of the lineplot, there are topoplots showing the scalp distribution of the GCMI values averaged in time windows of 150 ms. Statistically significant channels are denoted with a grey cross in the topoplots, while horizontal bars denote statistical significance.

**Fig. S2.**
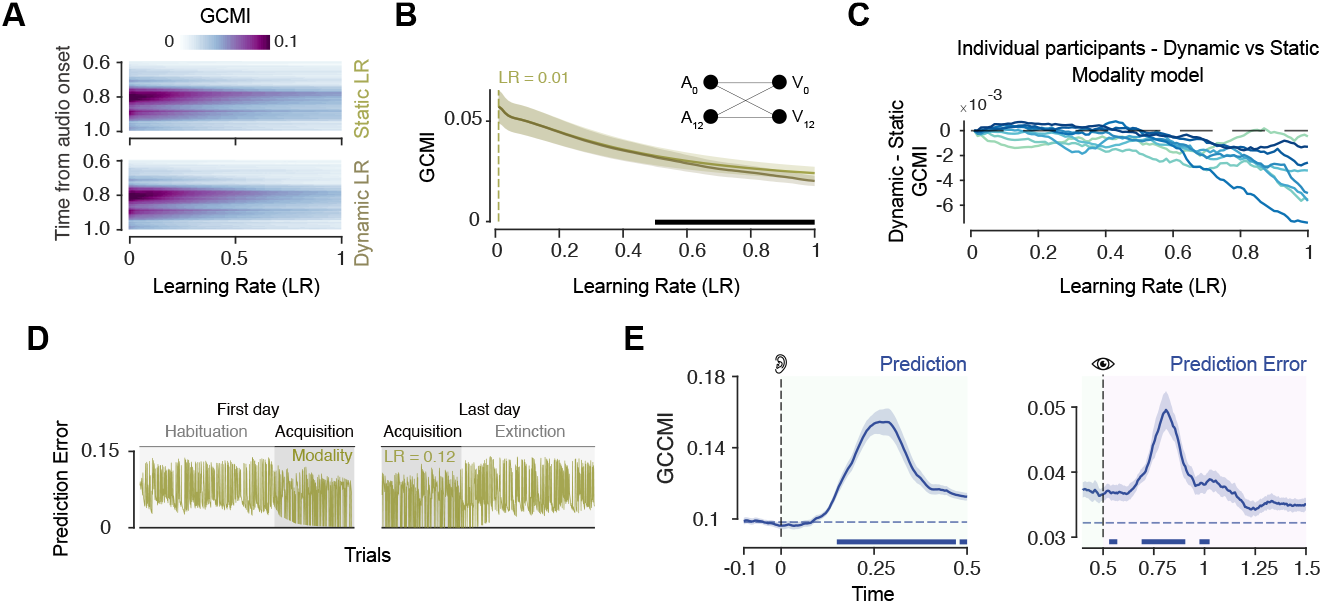
Model comparison between the stimulus-specific or sensory modality-level models. A. Group-average GCMI values for the Static and Dynamic learning rate (LR) models at the modality-level across the time window in the visual period (from 0.6 to 1 seconds from audio period onset) and the learning rate grid search parameter, ranging from 0.01 to 1. **B**. Lineplots showing GCMI averaged across the selected time window as a function of the learning rate. Shaded areas represent SEM across participants, while horizontal bars indicate statistical significance. **C**. Comparison between the Static and Dynamic LR Modality model at the participant level. Each line is a participant, showing the difference between the GCMI values of the two models as a function of the learning rate parameter. **D**. Prediction error trajectory (example from one participant) extracted from the Static LR modality model with the learning rate set to 0.12, similarly to the stimulus-specific model in the Fig. 4A. **E**. Conditional GCMI (GCCMI) values by fitting prediction and error trajectories of the stimulus-level model conditioned on the modality-level model’s trajectories (with LR=0.12, shown in **D**.). Shaded areas represent SEM across participants. Dotted lines indicate the onset of the audio and visual period and green and pink highlighted temporal regions represent the audio and visual period, respectively. Horizontal bars denote statistical significance.

**Fig. S3.**
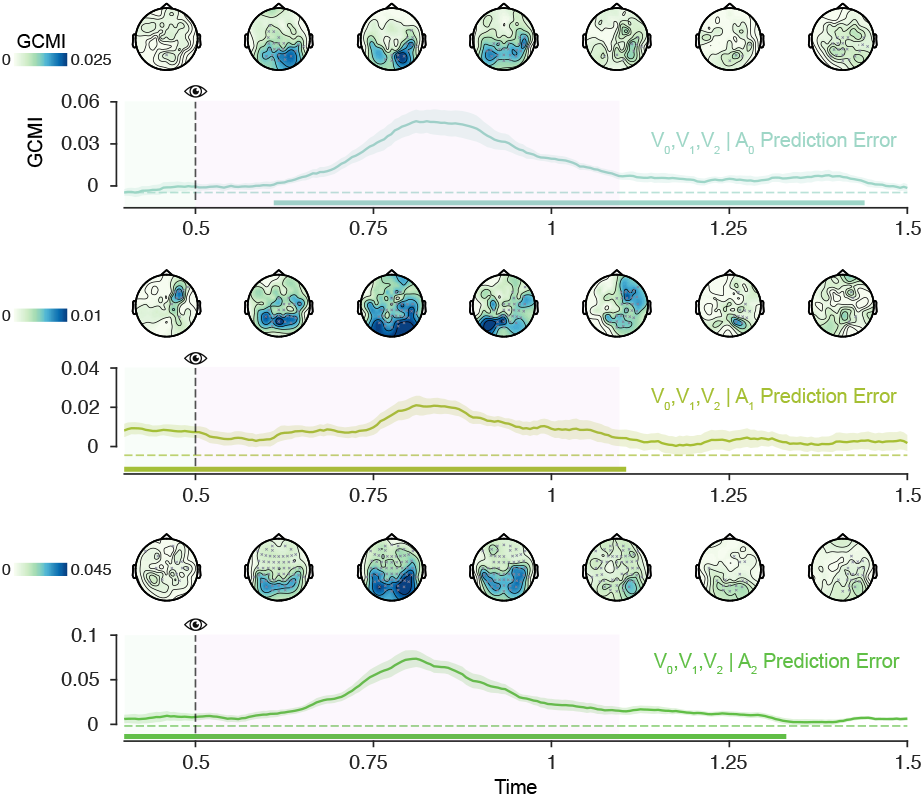
Cue-specific Prediction error encoding in the brain. Lineplots showing GCMI values for fitting the model predictions errors to the brain data (similarly to Fig. 4E but not conditioned on predictions). On top of each lineplot, topoplots show the scalp distribution of the GCMI values averaged in time windows of 150 ms. Statistically significant channels are denoted with a grey cross in each topoplot. Shaded areas represent SEM across participants. Dotted lines indicate the onset of the audio and visual period and green and pink highlighted temporal regions represent the audio and visual period, respectively. Horizontal bars denote statistical significance.

